# Adolescent Social Isolation Increases Vulnerability to Cocaine

**DOI:** 10.1101/320648

**Authors:** Anne Q. Fosnocht, Kelsey E. Lucerne, Alexandra S. Ellis, Nicholas A. Olimpo, Lisa A. Briand

## Abstract

Childhood and adolescent adversity is associated with a wide range of psychiatric disorders, including an increased risk for substance abuse. Despite this, the molecular mechanisms underlying how chronic stress during adolescence alters reward signaling remains largely unexplored. Understanding how adolescent stress increases addiction-like phenotypes could inform the development of targeted interventions both before and after drug use. The current study examined how adolescent-onset isolation stress affected behavioral, molecular, and physiological responses to cocaine in male and female mice. Adolescent-onset social isolation did not alter the ability of mice to learn an operant response for food, nor influence food self-administration or motivation for food on a progressive ratio schedule. However, male and female socially stressed mice exhibited an increase in motivation for cocaine and cocaine seeking during a cue-induced reinstatement session. Additionally, we demonstrated that adolescent-onset social isolation increased cocaine-induced neuronal activation, as assessed by Fos expression, within the nucleus accumbens core and shell, ventral pallidum, dorsal bed nucleus of the stria terminalis, lateral septum and basolateral amygdala. Taken together, the present studies demonstrate that social stress during adolescence augments the behavioral responses to cocaine during adulthood and alters the responsiveness of reward-related brain circuitry.

## 1. Introduction

Quality social interactions are fundamental to the mental health and well-being of an individual. A lack of social support during adolescence, either due to parental neglect, excessive bullying, or social exclusion, can increase the possibility of developing a psychiatric illness. These negative social experiences during adolescence increase the likelihood of alcohol and substance use during adolescence (Dube et al., 2006; Nelson et al., 1995; Somaini et al., 2011; Tharp-Taylor et al., 2009; Topper et al., 2011). Along with increased use and abuse during adolescence, adolescent stress also increases the likelihood of substance use and addiction in adulthood (Anda et al., 2002; Brents et al., 2015; Dube et al., 2002; Dube et al., 2003; Hoffmann et al., 2000; Kaltiala-Heino et al., 2000). However, as adolescent stress is often comorbid with other risk factors, unraveling a potential causal role for social stress in humans is difficult.

Preclinical models have shown that adolescent stress exposure can directly strengthen the effects of stimulants in adulthood. Adolescent stress exposure leads to sensitized responses to acute administration of drugs, strengthened contextual drug associations, increased escalation of cocaine taking on an extended access schedule, and increased motivation for cocaine on a progressive ratio schedule (Burke et al., 2013; Burke and Miczek, 2015; Burke et al., 2011; Cruz et al., 2012; Lepsch et al., 2005; Peleg-Raibstein and Feldon, 2011; Rodriguez-Arias et al., 2015). All of these studies utilized chronic physical stressors such as social defeat, restraint, and/or forced swim. There is also evidence that emotional stress during adolescence can lead to alterations in cocaine addiction phenotypes. Social isolation rearing in rats leads to increased rates of acquisition of cocaine self-administration and increased motivation for cocaine (Baarendse et al., 2014; Bozarth et al., 1989; Schenk et al., 1987). However, the effects of social isolation rearing without the reintroduction of peers on reinstatement of cocaine seeking are unclear.

Therefore, in these studies we first established that social isolation rearing increases motivation for cocaine and reinstatement of cocaine seeking in both male and female mice. Additional studies examined how adolescent stress interacts with chronic cocaine self-administration to alter responses to cocaine. To achieve this, we examined neural activation patterns within the mesocorticolimbic system following an acute cocaine injection in mice that had a history of cocaine self-administration and adolescent stress exposure. Our results indicate stress-dependent alterations in drug-taking behavior, as well as associated changes in neural activation.

## 2. Materials and Methods

### 2.1 Animals

This study used male and female c57/BL6 mice, exposed to social isolation beginning PND 21 through adulthood. Mice were randomly assigned isolation or group housing (2-5 animals per cage) at weaning (PND 21), then were all individually housed at the start of operant training (PND 60-80). All animals were housed in a temperature and humidity controlled animal care facility with a 12h light/dark cycle (lights on at 7:00 A.M.) and provided with cotton nestlets for enrichment. The Temple University Animal Care and Use Committee approved all procedures.

### 2.2 Drugs

Cocaine was obtained from the National Institutes of Drug Abuse Drug Supply Program (Bethesda, MD) and dissolved in sterile 0.9% saline.

### 2.3 Operant Food Training

At 8 weeks of age, mice were single-housed, food restricted to approximately 90% of their free feeding weight, and began operant food training. The animals were first trained to exhibit an operant response for sucrose pellets. They were placed in operant chambers (Med-Associates) and learned to spin a wheel manipulandum to receive a sucrose pellet. A white light, located over the active wheel, and 10s tone cue simultaneously occurred with administration of a pellet, followed by an 8s time-out with the house light off and no programmed consequences to wheel spins. Mice were food trained for 5 days on a fixed ratio schedule, where one wheel spin received a sucrose reward (FR1). After meeting criteria, they underwent 5 days of food training on a fixed ratio schedule where 5 wheel spins received a sucrose reward (FR5). During the first two sessions, the inactive wheel was immobilized. After that time, spinning on the inactive wheel has no programmed consequence. The mice were limited to a maximum of 50 pellets during the 60 min operant session. Following 10 days of food training, mice were tested for 1 day on a progressive ratio (PR) schedule, where the response requirement for each infusion increased until the subject did not fulfill the requirement. The response requirement was defined as R(i)=[5e^0.2i^-5] and the session ended if the animal took longer than 30 minutes to meet the requirement. The breakpoint is defined as the total number of responses or the final ratio completed.

### 2.4 Jugular Catheterization Surgery

Before surgery, mice were anesthetized with 80 mg/kg ketamine and 12 mg/kg xylazine. An indwelling silastic catheter was inserted into the right jugular vein and sutured in place. The catheter was threaded subcutaneously over the shoulder blade and routed to a mesh back mount platform (Strategic Applications, Inc) to secure the placement. Following surgery, catheters were flushed daily with 0.1 mL of antibiotic (Timentin, 0.93 mg/ml) dissolved in heparinized saline and sealed with plastic obturators while not in use.

### 2.5 Cocaine Self-administration

Mice were given 3-4 d to recover from surgery before the initiation of behavioral testing. The cocaine self-administration behavior was tested in 2-hour sessions (6 d per week) in the same chamber used for sucrose pellet self-administration. However, wheel responding now delivered an intravenous cocaine infusion (0.6 mg/kg/infusion), paired with the same compound cue, under the same fixed ratio(FR1) schedule as food training. After 10 days of a fixed ratio reward schedule, the mice were tested for 1 day on a progressive ratio (PR) schedule, where the response requirement for each infusion increased until the subject did not fulfill the requirement. The response requirement was defined as R(i)=[5e^0.2i^-5] and the session ended if the animal took longer than 30 minutes to meet the requirement. The breakpoint is defined as the total number of responses or the final ratio completed. Following PR, cocaine-seeking behavior was extinguished by replacing cocaine with 0.9% saline and removing the light and tone cues, previously paired with cocaine delivery. Daily 2 h extinction sessions occurred until animals performed <25% of their self-administration responding (average of last 3 days). Twenty-four hours after meeting this extinction criterion, animals underwent a cue-induced reinstatement session. The light and tone cues were presented non-contingently for 20 seconds, every 2 minutes during the first 10 minutes of the session. For the remainder of the session, the cues were presented contingent with operant responding, as they were during the cocaine self-administration phase. During the reinstatement session however, animals received saline infusions following responses on the active wheel. Catheter patency was tested once a week and animals were removed from the study if found to have lost patency.

### 2.6 Tissue Extraction

In order to examine the effect of isolation stress on the cocaine response in the brain, a separate group was exposed to the adolescent isolation, along with group-housed controls, and at 8 weeks of age underwent operant food training, jugular catheterization, and 10 days of cocaine self-administration as described above. Twenty-four hours after their last self-administration session, mice were either given an acute injection of cocaine (15 mg/kg) or taken from their home cage. 30 minutes following the cocaine injection, the mice received 100mg/kg pentobarbital (i.p.) before perfusion with 60 mL ice-cold PBS followed by 60 mL 4% PFA dissolved in ice-cold PBS. Brains were extracted and put in 4% PFA for 24 h before storage in 30% sucrose dissolved in PBS with 1% sodium azide. Coronal sections (40 μm) were taken using a cryostat and used for immunohistochemical staining.

### 2.7 Immunohistochemistry

We rinsed sections in 0.5M TBS 3 times for 5 minutes each then quenched with 0.30% hydrogen peroxide for 15 minutes. Next, tissue was blocked in 10% DS-TBS for 1 h before incubation in Fos primary antibody (Santa Cruz H-125) for 2 nights. On day 3, sections were rinsed again in .1%TBS-TX 3 times before a 2 h incubation in biotin donkey anti-rabbit. Following another rinse in TBS-TX, the sections were incubated in S-HRP (Vectastain) for 1 h then rinsed in TBS-TX 3 times. Finally, the tissue was incubated in DAB-Nickel for 5-10 minutes before the final rise in TBS until slide mounting.

### 2.8 Cell Counting

Fos-immunolabeling was bilaterally quantified from at least 2 sections per mouse and averaged to determine the profile of each brain region. The experimenter quantifying was blind to group assignments. Anatomical regions were determined by the stereotaxic atlas of Franklin and Paxinos (Franklin and Paxinos, 2013). Fos images were quantified using ImageJ software.

### 2.10 Statistical Analysis

All self-administration experiments were analyzed with two-way ANOVAs and post hoc comparisons as noted. The number of Fos immuno-labeled cells in the brain regions of interest was determined for each animal. Group differences in Fos immunoreactivity were analyzed using two-way ANOVAs with condition (isolation or group housing) and cocaine experience as the independent variables and number of cells as the dependent variable. Sidak’s post hoc comparisons were made when main effects or interactions were detected (p<0.05).

## 3. Results

### 3.1 Adolescent-onset social isolation in male mice increases responding for cocaine on an FR1 schedule of reinforcement

Mice socially isolated at weaning and group-housed controls were trained for ten days to learn the operant task for a food reward. Adolescent-onset social isolation did not alter the ability of male or female mice to acquire an operant response for food, nor alter their food intake or responding (Fig. 1; *males:* effect of session, pellets= *F*(9,396)=2.035, p=0.034; responses= *F*(9,396)=6.0, p<0.0001; *females:* effect of session, pellets= *F*(9,297)=3.02, p=0.0018; responses= *F*(9,297)=5.2, p<0.0001). After ten days of food training, all the mice underwent jugular catheterization surgery and completed ten days of cocaine self-administration. Adolescent-onset social isolation did not alter cocaine intake in male or female mice (Fig. 2a,b; *males:* effect of session, *F*(9,297)=3.11, p=0.0013, effect of stress, *F*(9,288)=1.0, p=0.33; *females:* effect of session, F(9,189)=0.89, p=0.54, effect of stress, F(9,189)=0.99, p=0.33). Following adolescent-onset social isolation, male mice exhibited an increase in responding for cocaine on an FR1 schedule, but female mice did not (Fig. 2c,d; *males:* effect of stress, *F*(9,297)=4.21, p=0.048; *females:* effect of stress, *F*(9,189)=0.63, p=0.44).

**Figure 1.**
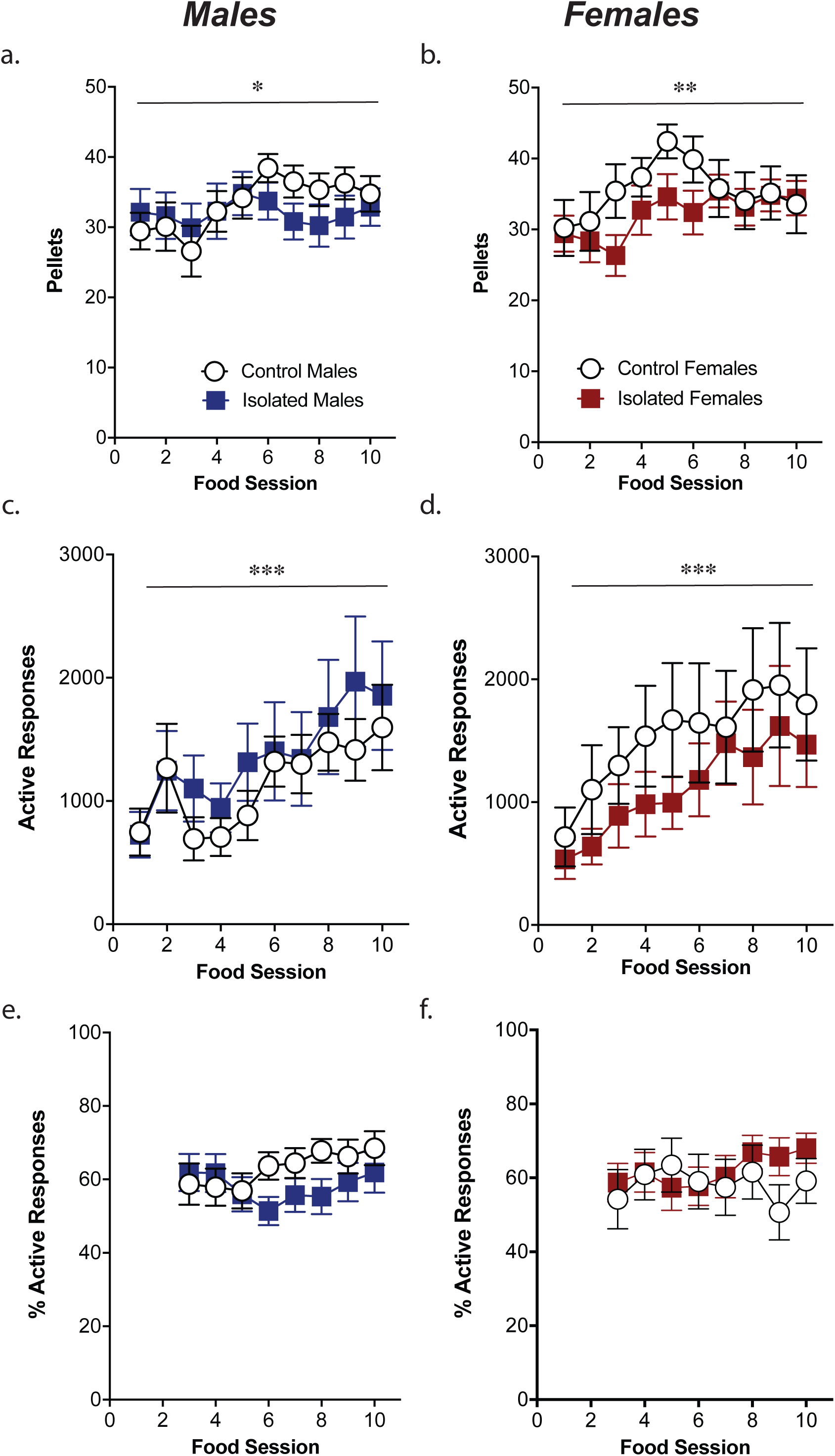
Adolescent isolation stress does not affect operant learning in food self-administration. Male and female control and stress exposed mice showed consistent levels of pellets earned (a,b), a gradual increase in active responses (c,d), and equal levels of percent active responding over 10 days of food training (e,f). *p<.05, **p<.01, **p<.001 main effect of session, n=14–20/group

**Figure 2.**
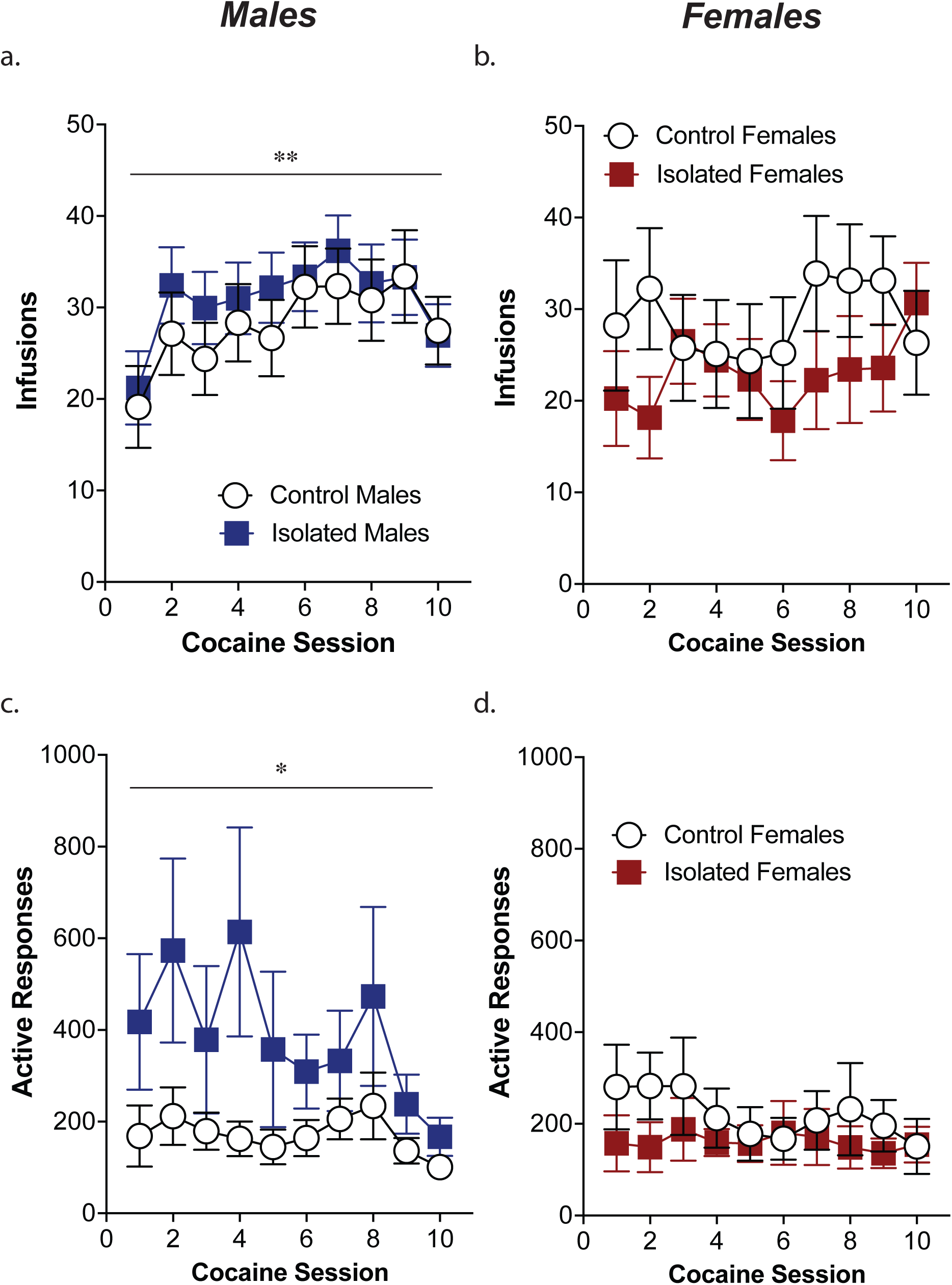
Adolescent isolation stress increases responding for cocaine on an FR1 schedule in males but not females. Male and female control and stress exposed mice earned equal numbers of infusions over the course of 10 days of cocaine self-administration (a,b; n=16–19/group). However, male mice exposed to adolescent-onset social isolation exhibit higher rates of responding for cocaine compared to male controls (c). Female mice isolated during adolescence do not exhibit any differences in responding compared to controls (d; n=11–13/group). **p<.01 main effect of session; *p<.05 main effect of stress

### 3.2 Adolescent-onset social isolation motivation for cocaine and potentiates cue-induced reinstatement of drug seeking in both male and female mice

However, isolation stress led to an increase in responding on a progressive ratio schedule for cocaine in both male and female mice (Fig. 3a; effect of stress, *F*(1,50)=4.14, p=0.047; effect of sex, *F*(1,50)=0.24, p=0.63). Isolation stress did not alter extinction learning, as male and female stressed and control mice exhibited similar rates of extinction (Fig. 3b; effect of stress, *F*(1,37)=0.086, p=0.77; effect of sex, *F*(1,37)=0.054, p=0.82) and similar levels of responding on the final day of extinction (Fig. 3c; effect of stress, *F*(1,37)=2.47, p=0.13; effect of sex, *F*(1,37)=2.5, p=0.12). Twenty-four hours after the last extinction session, mice were exposed to a cue-induced reinstatement session in which active responses resulted in presentation of the previously drug-paired cues, in the absence of the drug. Both stress and control groups showed significant reinstatement during this session, however, the exposure of isolation stress during adolescence caused a potentiated reinstatement of drug seeking in both male and female mice (Fig. 3d; effect of stress, *F*(1,37)=5.46, p=0.025; effect of sex, *F*(1,37)=1.52, p=0.23).

**Figure 3.**
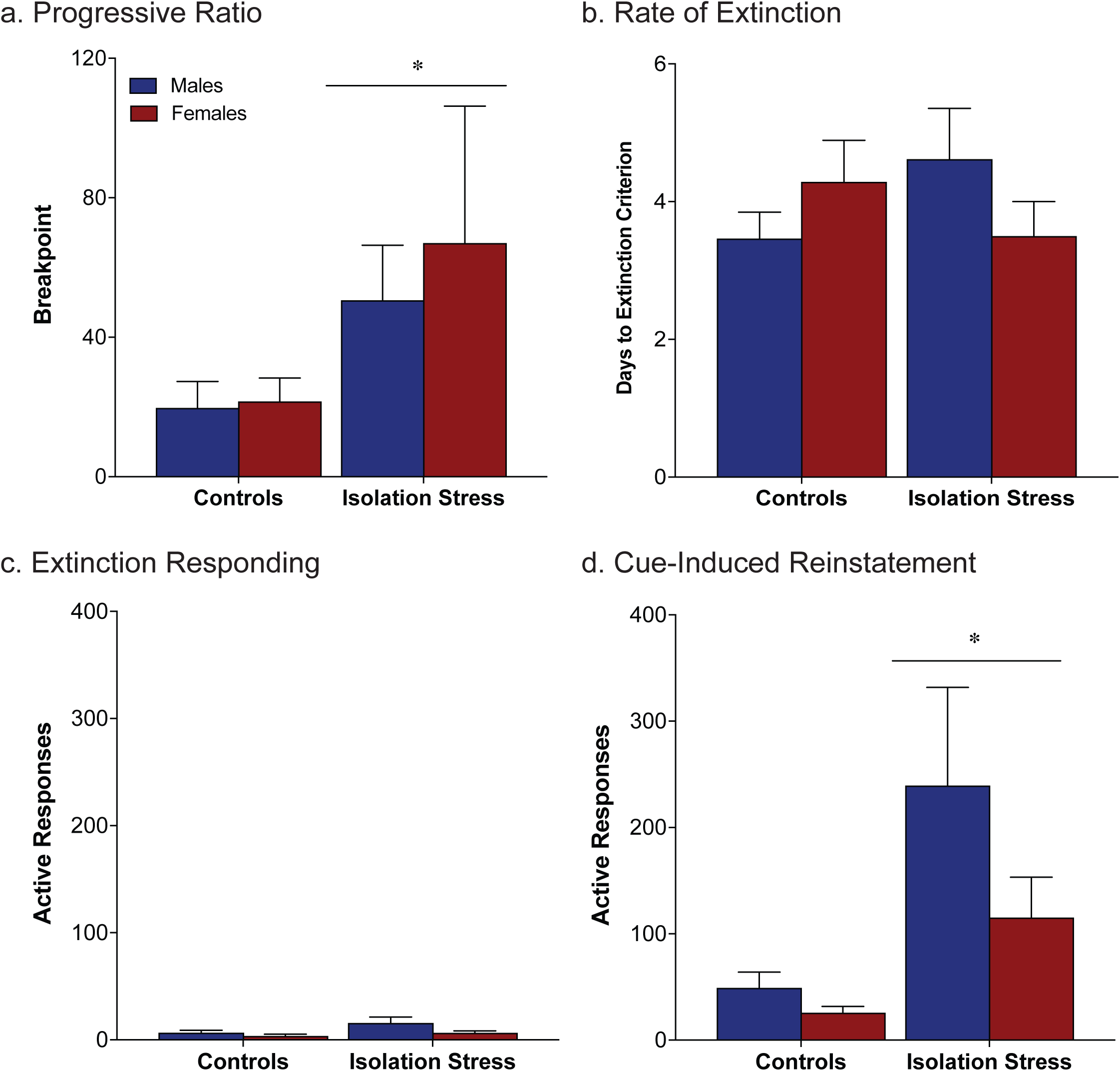
Isolation stress during adolescence increases motivation for cocaine and cue-induced cocaine seeking in adulthood. Male and female adolescent stress exposed mice respond more on a progressive ratio schedule of reinforcement, exhibiting a higher breakpoint (a; n=10–18/group). No differences were seen between the groups in the rate of extinction (b) or the responding on the final day of extinction (c). Twenty-four hours after the final extinction session, during the cue-induced reinstatement session male and female mice exposed to adolescent social isolation exhibited higher rates of responding compared to controls (d; n=8–12/group). *p<.05 main effect of stress

### 3.3 Cocaine-induced c-fos expression is increased following adolescent-onset social isolation stress and chronic cocaine self-administration

To better understand the neurobiological mechanisms underlying the behavioral differences observed following adolescent-onset social isolation, we looked at cocaine-evoked c-Fos expression in several brain regions following chronic cocaine exposure and adolescent stress. Due to the similar trends seen in the motivation for cocaine and the reinstatement of cocaine seeking, male and female mice were grouped together for these studies. Mice underwent operant food training, catheter implantation, and 10 days of cocaine self-administration. Twenty-four hours after their last self-administration session, mice were either given an acute injection of cocaine (15 mg/kg) or taken from their home cage and brains were collected 30 minutes later for immunohistochemical staining. Adolescent-onset isolation coupled with chronic cocaine exposure causes a significant increase in cocaine-induced c-Fos expression in the nucleus accumbens core and shell, dorsal bed nucleus of the stria terminalis (BNST), lateral septum, basolateral amygdala and a trend towards an increase in the ventral pallidum (Fig. 4).

**Figure 4.**
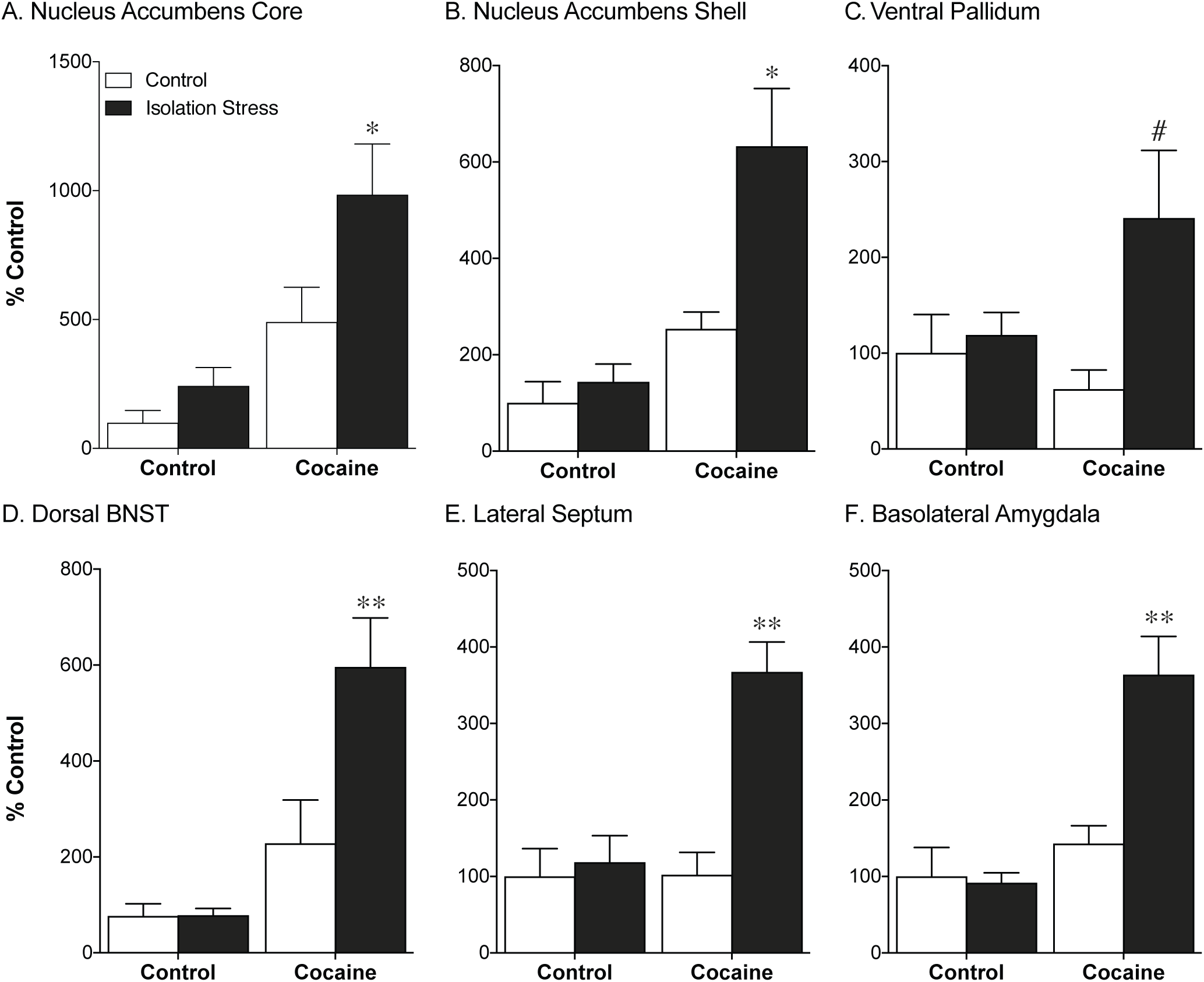
Chronic cocaine exposure after adolescent stress causes an increased fos response in the nucleus accumbens core and shell, basolateral amygdala, and lateral septum. Stress exposed mice show significantly greater Fos activation in the nucleus accumbens core and shell after chronic cocaine compared to group housed controls (a,b). Similarly, chronic cocaine in stress exposed mice causes a potentiated Fos response in the basolateral amygdala and lateral septum (c,d). #p=.05, *p<.05, **p<.01 post-hoc control cocaine vs. isolation cocaine; n=5–9/group

## 4. Discussion

Here we present evidence that social isolation beginning in adolescence leads to increased cocaine taking and seeking in adulthood. This increase in behavioral response to cocaine is accompanied by an increase in cocaine-induced c-Fos expression within regions implicated in reward processing as well as regions that mediate stress responsivity. Furthermore, adolescent social isolation leads to increased AMPA transmission within the nucleus accumbens. We suggest that this increased behavioral responsivity to cocaine may be mediated by heightened glutamate activity within reward and stress related circuitry.

### 4.1 Adolescent social isolation promotes vulnerability to addiction-like behaviors in adulthood in both male and female mice

Early life stress is a risk factor for the development of addiction in humans (Brents et al., 2015; Dube et al., 2003; Dube et al., 2006). Social stressors are highly prevalent and among the most powerful stressful stimuli experienced by humans (Backstrom and Winberg, 2013). Modeling this increased risk in animals can provide us with a means to identify neural mechanisms that underlie increased vulnerability in humans. Rodents who are socially isolated at weaning exhibit enhanced alcohol consumption (Chappell et al., 2013; McCool and Chappell, 2009), enhanced cocaine intake (Ding et al., 2005), enhanced locomotor response to amphetamine (Herrmann et al., 2014), and enhanced cocaine-evoked dopamine release (Yorgason et al., 2016). In the present study, social isolation at weaning led to an increase in cocaine taking during self-administration, increased motivation for cocaine on a progressive ratio schedule, and increased drug seeking during cue-induced reinstatement. While the pathologically relevant time period for isolation is adolescence, the socially isolated animals in the present study were not reintroduced to group housing conditions before beginning the operant training paradigm. Instead, animals in the control group were isolated at this time due to issues with group housing animals with indwelling catheters. While it is possible that the duration of the social isolation influenced the current findings, the control animals had been socially isolated for 3 weeks at start of the cocaine self-administration phase, the enhanced addictive-like phenotypes were unique to animals isolated during adolescence. Moreover, the effects of adolescent social isolation were specific to cocaine as no differences in food intake or motivation for food were seen.

These effects are consistent with what has been seen following other forms of adolescent stress. In particular, numerous studies have demonstrated that adolescent social defeat increases addictive-like phenotypes in male rodents. Specifically, increased conditioned place preference to cocaine and MDMA is seen following adolescent social defeat stress (Garcia-Pardo et al., 2015; Montagud-Romero et al., 2017). Similarly, both adolescent social defeat and social isolation increase cocaine self-administration in male rats (Burke and Miczek, 2015; Yajie et al., 2005). However, post-weaning social isolation served to protect rats from this increase in cocaine taking seen following social defeat in pair-housed animals (Burke and Miczek, 2015). In contrast to the increase in addictive phenotypes seen following adolescent social stress, stress earlier in life may be protective. Maternal separation stress leads to decreased cocaine self-administration, particularly in males (Martini and Valverde, 2012; Matthews et al., 1999; O'Connor et al., 2015). This suggests that perhaps adolescence is a critical period for stress exposure to predispose individuals to develop addiction.

The majority of models examining social stress have utilized only male animals due to male’s innate territorial aggression toward other males, a behavior that is absent in females under normal conditions (Haller et al., 1999; Takahashi et al., 2017). The current study is the first to demonstrate that adolescent social stress can increase addictive-like behaviors in adult females. As human females are more likely to develop stress-related psychiatric disorders (Bangasser and Valentino, 2012), developing a model to examine the influence of adolescent social stress on addictive-like behaviors in females is critical.

### 4.2 Adolescent social isolation increases the neuronal response to cocaine in adulthood

As the brain is still developing throughout childhood and adolescence, stress during this period of life may alter the development of reward circuitry. Human functional imaging studies indicate that childhood trauma leads to dysfunction in reward processing in adulthood (Dillon et al., 2009; Mehta et al., 2010). Furthermore, PET binding studies suggest that adults with a history of childhood trauma exhibit increased dopamine response to stimulants (Oswald et al., 2014). However, it is not clear from these studies whether these alterations in reward processing are a consequence of stress exposure or represent some common vulnerability to stress and addiction that precedes trauma. The current study determined that adolescent social isolation increases cocaine-induced activation of stress and reward circuitry. In group-housed animals with a history of cocaine self-administration, acute cocaine exposure led to an increase in c-Fos expression in the nucleus accumbens core and shell and dorsal bed nucleus of the stria terminalis. However, in socially isolated mice with a history of cocaine self-administration, acute cocaine increased c-Fos to a greater extent in these three regions as well as engaging additional brain regions: ventral pallidum (VP), lateral septum (LS), and the basolateral amygdala (BLA).

The ventral pallidum receives input from the nucleus accumbens and serves to integrate information from multiple cell populations. Chronic cocaine exposure leads to potentiated output in accumbal MSNs (Creed et al., 2016). Therefore, the increase in c-Fos activation within the accumbens and VP could represent a further potentiation of this response by adolescent stress exposure. Future work could examine the ability of stress to influence plasticity within this circuit. Cocaine also led to activation of stress-related brain circuits in animals that received adolescent social stress. The BLA projections to both the nucleus accumbens and the orbitofrontal cortex play key roles in cue-induced cocaine seeking (Arguello et al., 2017; Lasseter et al., 2011; Stefanik and Kalivas, 2013). Furthermore, early life stress leads to increased spine density and synaptic excitability within the BLA (Guadagno et al., 2018). Coupled with our findings, this supports that increased activation within the BLA could mediate the increase in cue-induced cocaine seeking seen following adolescent social isolation. Recently, the LS has also been implicated in cocaine seeking behavior. Inputs from the LS to the ventral tegmental area are activated during cue-induced reinstatement (Mahler and Aston-Jones, 2012) and projections from the dorsal hippocampus to the LS are activated during context-induced reinstatement (McGlinchey and Aston-Jones, 2017). Further examination of the influence of adolescent stress on inputs both to and from the LS could provide insight into the behavioral differences seen in reinstatement behavior in the current study.

Our results are consistent with previous work demonstrating that isolation rearing increases the ability of an acute cocaine injection to induce Fos expression within the nucleus accumbens core and shell (Howes et al., 2000). The current study expands this to demonstrate cocaine-induced changes in Fos expression in animals with a history of cocaine self-administration. We see more widespread increases in regions implicated in stress along with addiction. Previous studies have demonstrated increases in Fos expression following acute cocaine in regions where we did not see increases in the group housed animals. This includes studies done in drug naïve mice demonstrating increased Fos expression the lateral septum (Briand et al., 2010) and studies examining Fos expression immediately following self-administration that have seen increased Fos in the BLA and VP (Zahm et al., 2010). There may be something unique about the combination of self-administration and experimenter administered cocaine or these differences may be due to across species differences as all these studies were done in rats. Additionally, a large body of work has examined Fos expression in cocaine-experienced mice following periods of abstinence or extinction. These paradigms lead to robust increases in Fos expression throughout brain regions implicated in reward, including the nucleus accumbens and regions implicated in stress reactivity, including the BLA (Hearing et al., 2008; Zavala et al., 2007; Zavala et al., 2008). These data, coupled with the reinstatement data in the current study, suggest that we might see a different pattern of Fos expression if we examined animals after reinstatement.

### 4.3 Conclusion

Taken together, the current results indicate that adolescent social isolation leads to an increase in cocaine self-administration, motivation for cocaine, and cocaine seeking during cue-induced reinstatement. These increases are specific to drug reward as social isolation did not impact food self-administration or motivation for food. Accompanying these increases in behavioral response to cocaine is an increase in the neuronal response to cocaine as evidenced by increased c-Fos expression. These alterations in cocaine-induced c-Fos may be mediated by differences in AMPA transmission within the nucleus accumbens. These results highlight the role of adolescent stress in addiction and suggest that individual vulnerabilities to addiction may be mediated by alterations in AMPA transmission in the nucleus accumbens.

## Contributors

The project idea was initiated and designed by LAB, in consultation with AQF. AQF, KEL, and ASE planned and executed the behavioral experiments. AQF, KEL, and NAO performed the immunohistochemistry experiments. AQF and LAB wrote the first draft of the manuscript and all authors contributed to the final text.

## Acknowledgements

This work was supported by National Institute on Drug Abuse (NIDA) Grant R00 DA033372 (L.A.B.) and a Brain & Behavior Research Foundation NARSAD award (L.A.B.).

We thank Dr. Debra Bangasser for providing the feedback on the manuscript.

---

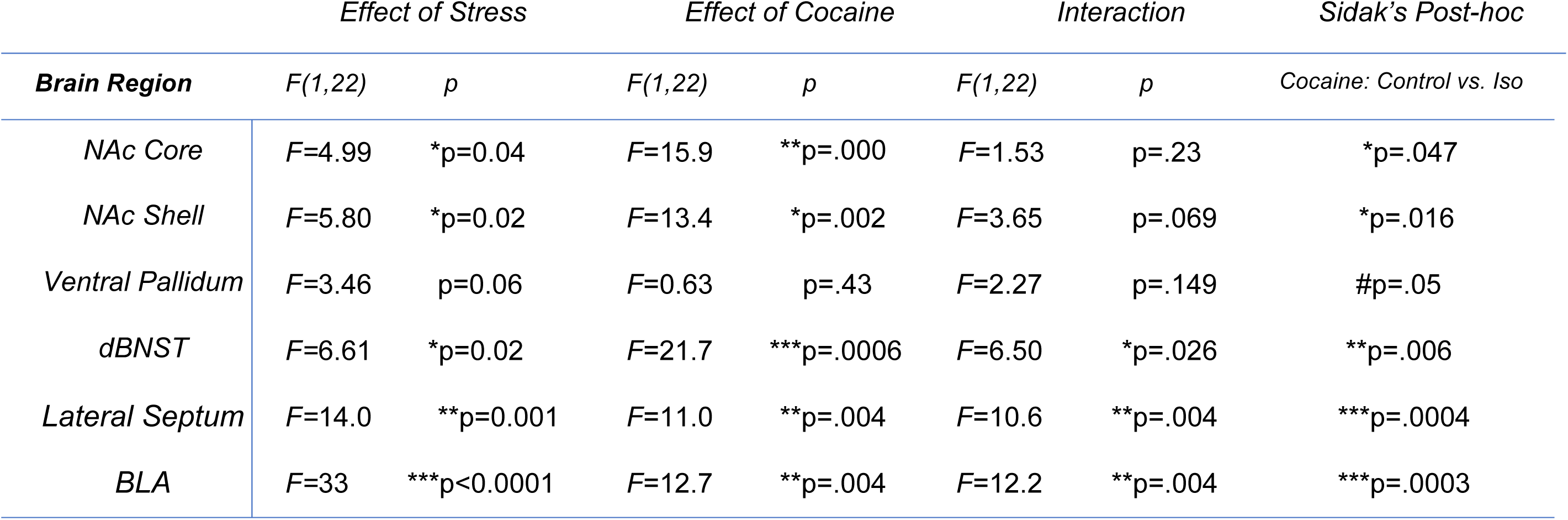

